# Combining Directed Evolution with Machine Learning Enables Accurate Genotype-to-Phenotype Predictions

**DOI:** 10.1101/2025.01.27.635131

**Authors:** Alexander J. Howard, Ellen Y. Rim, Oscar D. Garrett, Yejin Shim, James H. Notwell, Pamela C. Ronald

## Abstract

Linking sequence variation to phenotypic effects is critical for efficient exploitation of large genomic datasets. Here we present a novel approach combining directed evolution with protein language modeling to characterize naturally-evolved variants of a rice immune receptor. Using high-throughput directed evolution, we engineered the rice immune receptor *Pik-1* to bind and recognize the fungal proteins Avr-PikC and Avr-PikF, which evade detection by currently characterized *Pik-1* alleles. A protein language model was fine-tuned on this data to correlate sequence variation with ligand binding behavior. This modeling was then used to characterize *Pik-1* variants found in the 3,000 Rice Genomes Project dataset. Two variants scored highly for binding against Avr-PikC, and *in vitro* analyses confirmed their improved ligand binding over the wild-type *Pik-1* receptor. Overall, this machine learning approach identified promising sources of disease resistance in rice and shows potential utility for exploring the phenotypic variation of other proteins of interest.

## Main Text

Protein language models (PLMs) like ESM-2^1^ are transformer-based neural networks trained on enormous sets of evolutionarily-derived proteins to learn protein sequence, structure, and functional information. After training on this data, PLMs can be used to distill any input protein sequence into a high-dimensional numerical representation called an “embedding”. These embeddings have been used in the past as inputs for specialized machine learning tasks^2–5^.

Alternatively, PLMs themselves can be “fine-tuned” to produce a specialized model that directly predicts the properties of an input protein sequence^6,7^. Fine-tuning is a process which takes a pre-trained model, optionally adjusts the model architecture, and trains the entire model on a specialized dataset to predict the characteristics measured in that data. A major benefit of this approach is that backpropagation during training extends into the language model weights, adapting the entire model towards the prediction task^6,7^. Fine-tuned PLMs have been previously used to accurately predict the effect of missense mutations on enzyme function^8^, protein stability^9^, and protein-protein interactions^7,10^. This flexibility and accuracy makes fine-tuned PLMs a useful tool for predicting the effects of sequence variation on phenotypes of interest, which would be especially valuable for exploring large genomic datasets.

*Magnaporthe oryzae* is the fungal pathogen responsible for rice blast, a disease that can cause a yield loss of 10-30% in rice and destroys enough rice each year to feed 60 million people^11,12^. Several genes in rice can confer resistance against blast disease, including the immune receptor *Pik-1* which binds the *M. oryzae*-secreted protein Avr-Pik via an integrated heavy metal-associated (HMA) domain^13,14^. Several variants of Avr-Pik (Avr-PikA through Avr-PikF) have been identified across *Magnaporthe* strains, each featuring sequence variations that can weaken or break the HMA/ligand interactions required for *Pik-1* to initiate an immune response^15^ (**Fig. 1a**). Variations in the *Pik-1* HMA domain have a significant impact on the recognition profile of a given *Pik-1* receptor allele, as shown by the Pikp-1 and Pikh-1 alleles which differ by only one residue in the HMA domain but in turn recognize one Avr-Pik variant and four Avr-Pik variants, respectively^16^ (**Fig. 1a**). These differing activities *in planta* are linked to the binding affinity of the HMA domain to each Avr-Pik variant^14,16^. The importance of HMA/ligand binding for *Pik-1* functionality has been previously exploited to engineer *Pik-1* receptors with expanded Avr-Pik recognition profiles^17,18^. Previously, we outlined a method to engineer enhanced Pikh-1 HMA domain binding against both Avr-PikC and Avr-PikF^19^, which no natural allele of *Pik-1* has been shown to achieve (**Fig. 1b**). Millions of Pikh-1 HMA domain variants were generated with error-prone PCR and transformed into yeast cells for yeast-surface display (YSD). This starting YSD library contained ∼2 x 10^7^ variants, each featuring on average 2.1 amino acid substitutions along the 78 amino acid-long domain^19^. The variant library was then screened for binding against fluorescently-labeled Avr-PikC or Avr-PikF. Variants with enhanced binding against these ligands relative to the wild-type Pikh-1 HMA domain were selected via fluorescently-activated cell sorting (FACS) and sequenced.

**Fig. 1:**
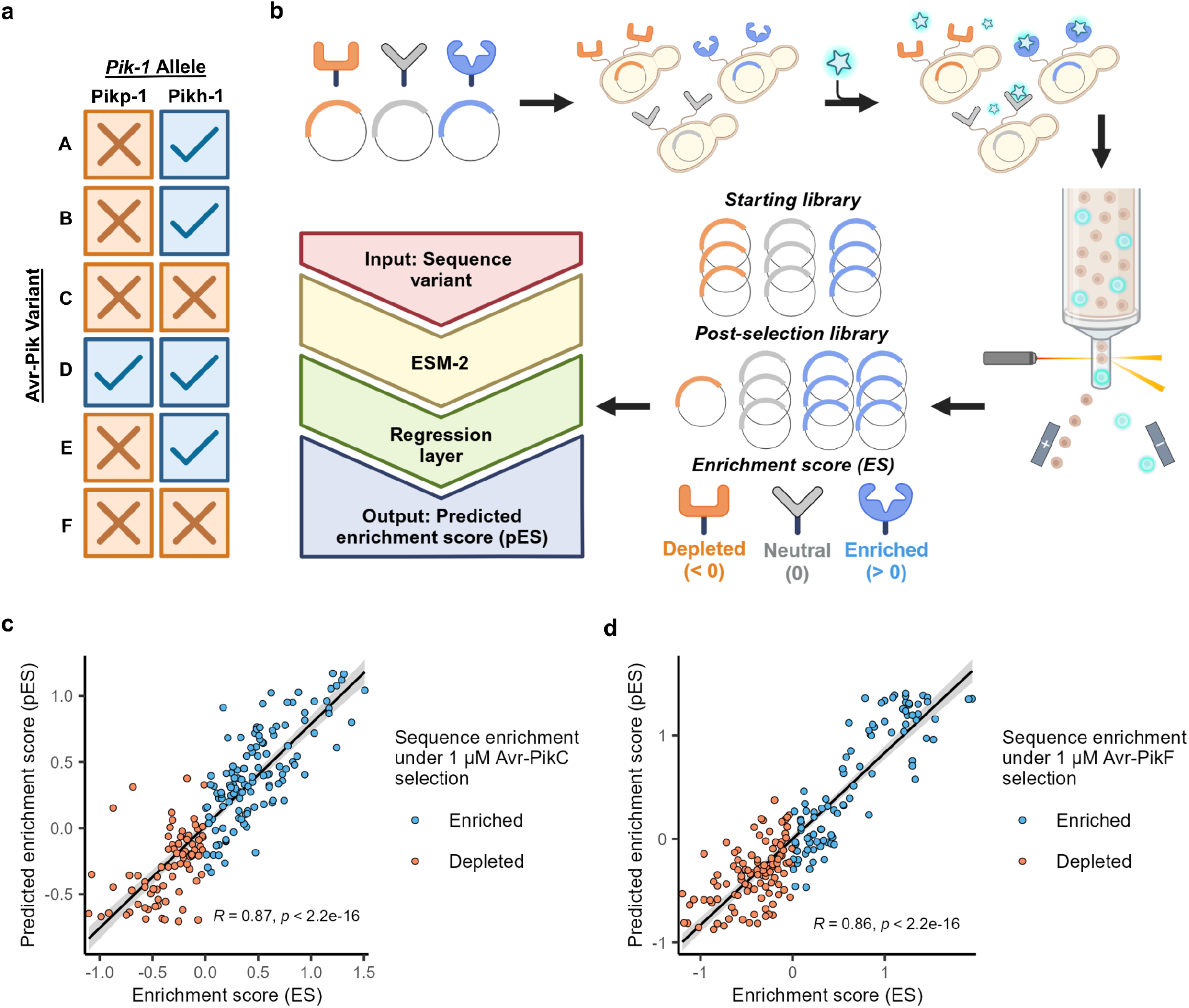
ESM-2 models fine-tuned on directed evolution data strongly correlate sequence variants with effects on ligand binding. **a**, The recognition profile of *Pik-1* alleles against different Avr-Pik variants is shown. Receptor/ligand combinations which trigger immune signaling are shown in blue while combinations which do not are shown in orange. **b**, Schematic of YSD directed evolution of the Pikh-1 HMA domain and fine-tuning of ESM-2 to predict variant performance. **c, d**, pES (y-axis) compared to true ES (x-axis) for validation sequences are shown for ESM-2 models fine-tuned on 1μM Avr-PikC (left) and 1μM Avr-PikF (right) selection data. Depleted sequences are shown in orange and enriched sequences are shown in blue. Line of best fit with 95% confidence interval and Spearman correlation coefficient (R) are also shown.

We used our directed evolution data to fine-tune ESM-2 to predict *Pik-1* HMA domain variant binding against Avr-PikC and Avr-PikF. Receptor performance was quantified with an enrichment score (ES), which measured the relative change in sequence abundance between the starting YSD library and post-selection YSD library. This data was split into training and validation sets to fine-tune ESM-2 and calculate a predicted enrichment score (pES) for input receptor variants. After training, the final Avr-PikC (**Fig. 1c**) and Avr-PikF (**Fig. 1d**) fine-tuned models both obtained a Spearman correlation coefficient R value over 0.85 on the validation data, indicating these models were able to strongly associate key sequence characteristics with changes in ligand binding. This modeling approach outperformed alternate models trained on ESM-2 embeddings, demonstrating the value of using fine-tuning to leverage PLM-distilled information (**Table S1, S2**). Given the robust performance of these fine-tuned models on our directed evolution data, we next tested their applicability towards phenotyping naturally-evolved *Pik-1* HMA domain variants for Avr-PikC and Avr-PikF binding.

Sequencing reads from the 3,000 Rice Genomes Project^20^ (3k RGP) were aligned against a reference genome to identify variants of the Pikh-1 HMA domain. 119 rice varieties returned full read coverage along the HMA domain, resulting in the identification of 13 unique HMA variants (**Fig. 2**), 11 of which to our knowledge had not been phenotypically characterized for binding against Avr-PikC or Avr-PikF. All sequence variants were input into our fine-tuned models to obtain pES values (**Fig. 2**). The Pikh-1 and Pikp-1 alleles were scored negatively for Avr-PikC and Avr-PikF binding, which aligned with previous phenotyping of these variants^16^. Ten variants received a positive pES value for Avr-PikC binding, and no variants were positively scored for Avr-PikF binding. Interestingly, our model which was trained on sequence data that lacked insertion or deletion mutations consistently scored *Pik-1* variants with an insertion in the middle of the HMA domain highly for Avr-PikC binding (**Fig. 2**). Given this observation, two variants with unique insertions and high pES values were chosen for downstream phenotyping: Vellai Kolomban (VK) and Sanhuangzhan-2 (SHZ-2).

**Fig. 2:**
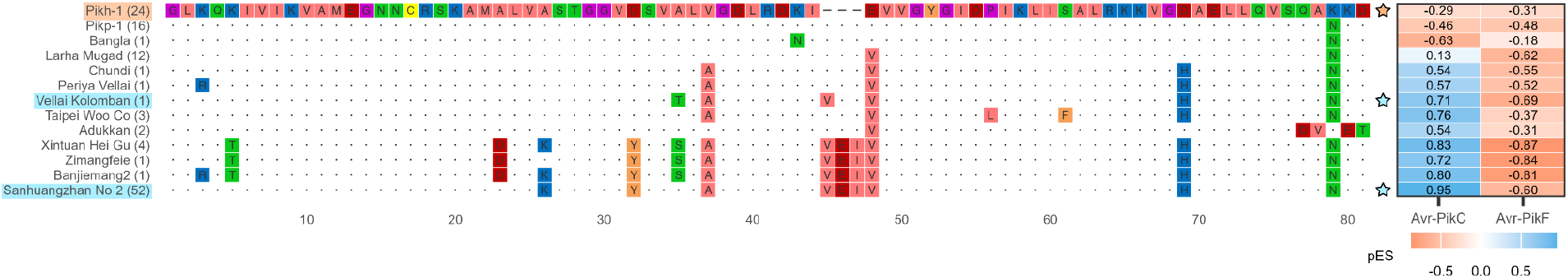
Several *Pik-1* alleles feature novel HMA domain variations predicted by fine-tuned ESM-2 to bind Avr-PikC. Multiple sequence alignment of *Pik-1* HMA domain variants identified from the 3k RGP dataset are shown, with residues differing from Pikh-1 (top row) shown in color. *Pik-1* variants without a known name are labeled with a representative rice variety carrying the allele. The number of occurrences of each variant is shown to the right of each variant name in parentheses. A table of pES values for Avr-PikC and Avr-PikF binding is shown to the right, with negative pES values in orange and positive pES values in blue. Variants selected for downstream testing are highlighted (left) and starred (right).

The Pikh-1, VK, and SHZ-2 *Pik-1* HMA domains were expressed with YSD and tested for binding against Avr-PikA, which is recognized by Pikh-1 and thus served as a positive control, and Avr-PikC at 1 μM concentration. These cells were imaged for ligand binding and sorted with FACS to quantitatively compare the binding behavior of each receptor (**Fig. 3a, 3b**). Pikh-1 showed the strongest binding against Avr-PikA, with VK and SHZ-2 both showing low to moderate interaction with the ligand. Pikh-1 showed minimal binding to Avr-PikC, with VK showing improved binding and SHZ-2 showing the highest affinity, closely following the predictions made by our fine-tuned model.

**Fig. 3:**
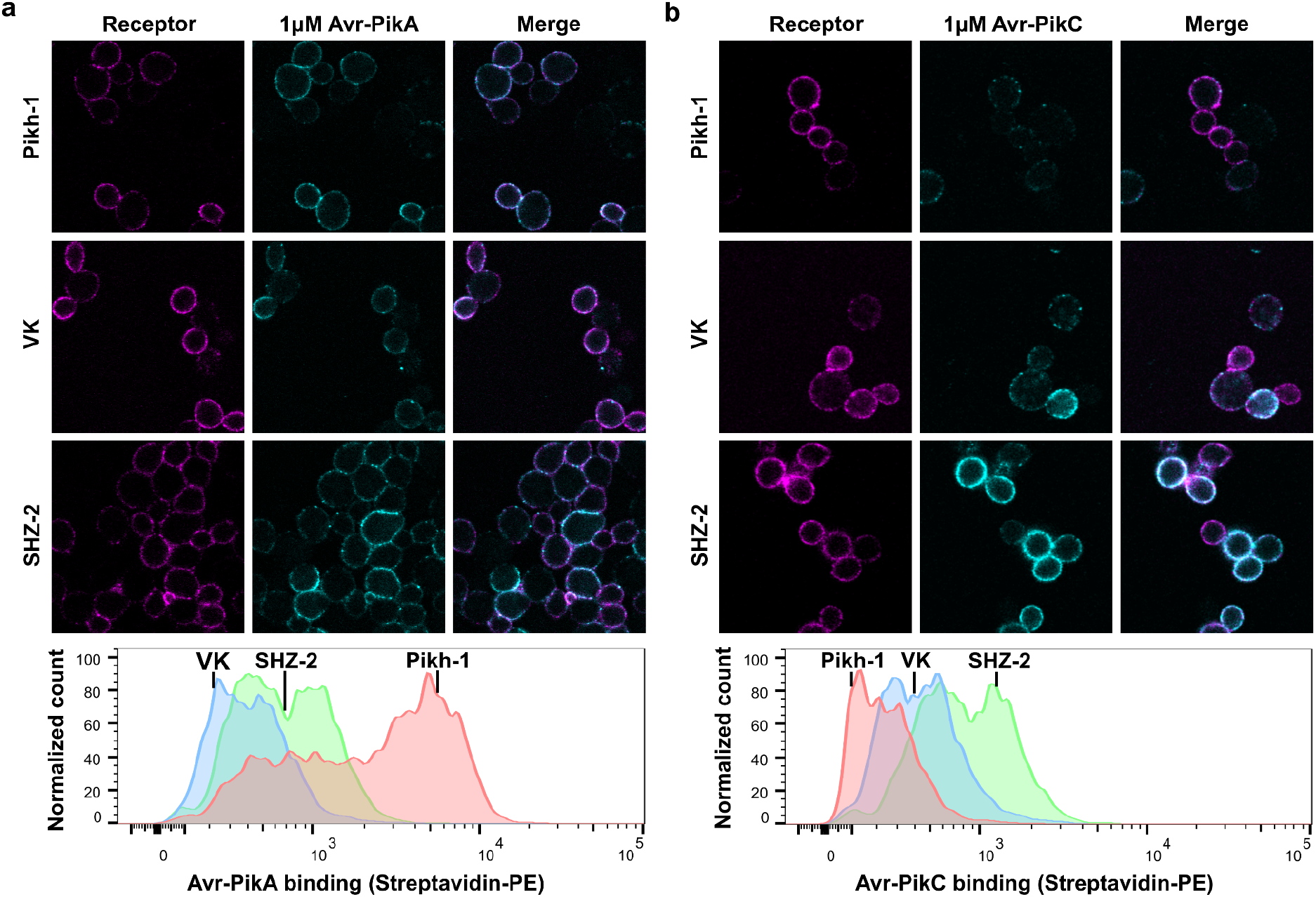
Fine-tuned ESM-2 predictions on *Pik-1* variant Avr-PikC binding are supported *in vitro*. **a, b**, Representative images of YSD clones expressing Pikh-1, VK, and SHZ-2 HMA domains binding to Avr-PikA (left) and Avr-PikC (right) at 1 μM are shown. Receptor expression is shown in magenta and ligand binding is shown in cyan. FACS measurements for individual YSD clones expressing Pikh-1 (red), VK (blue), or SHZ-2 (green) HMA domains against Avr-PikA (left) and Avr-PikC (right) at 1 μM are shown below.

To explore the generalizability of this approach, we searched for additional datasets which utilize protein mutagenesis to predict phenotypic effects. A mutagenesis scan of the human enzyme Nudix hydrolase 15 (NUDT15) by Suiter *et al*.^21^ was chosen to be modeled, as loss-of-function variations in this gene have been found to increase the risk of cytotoxicity in patients treated with thiopurine drugs^21–23^. Thiopurines are a frequently-used treatment for patients with leukemia and inflammatory bowel disease^24–27^, so accurately correlating NUDT15 sequence variation with cytotoxicity risk is crucial for optimizing patient treatment approaches. NUDT15 variant stability and functionality measurements made by Suiter *et al*. were used to create a functionality score (FS) for each variant, where positive scores indicated the enzyme retained functionality while negative scores indicated a loss of enzyme functionality. Any sequences that would be tested later in downstream phenotyping were filtered from the dataset, and all remaining variants were split into training and validation sets to fine-tune ESM-2 and calculate a predicted functionality score (pFS) for input NUDT15 variants (**Fig. 4a**). The final fine-tuned model obtained an R value of 0.76 on the validation data, indicating the model was able to effectively associate NUDT15 sequence variations with changes in enzyme functionality.

**Fig. 4:**
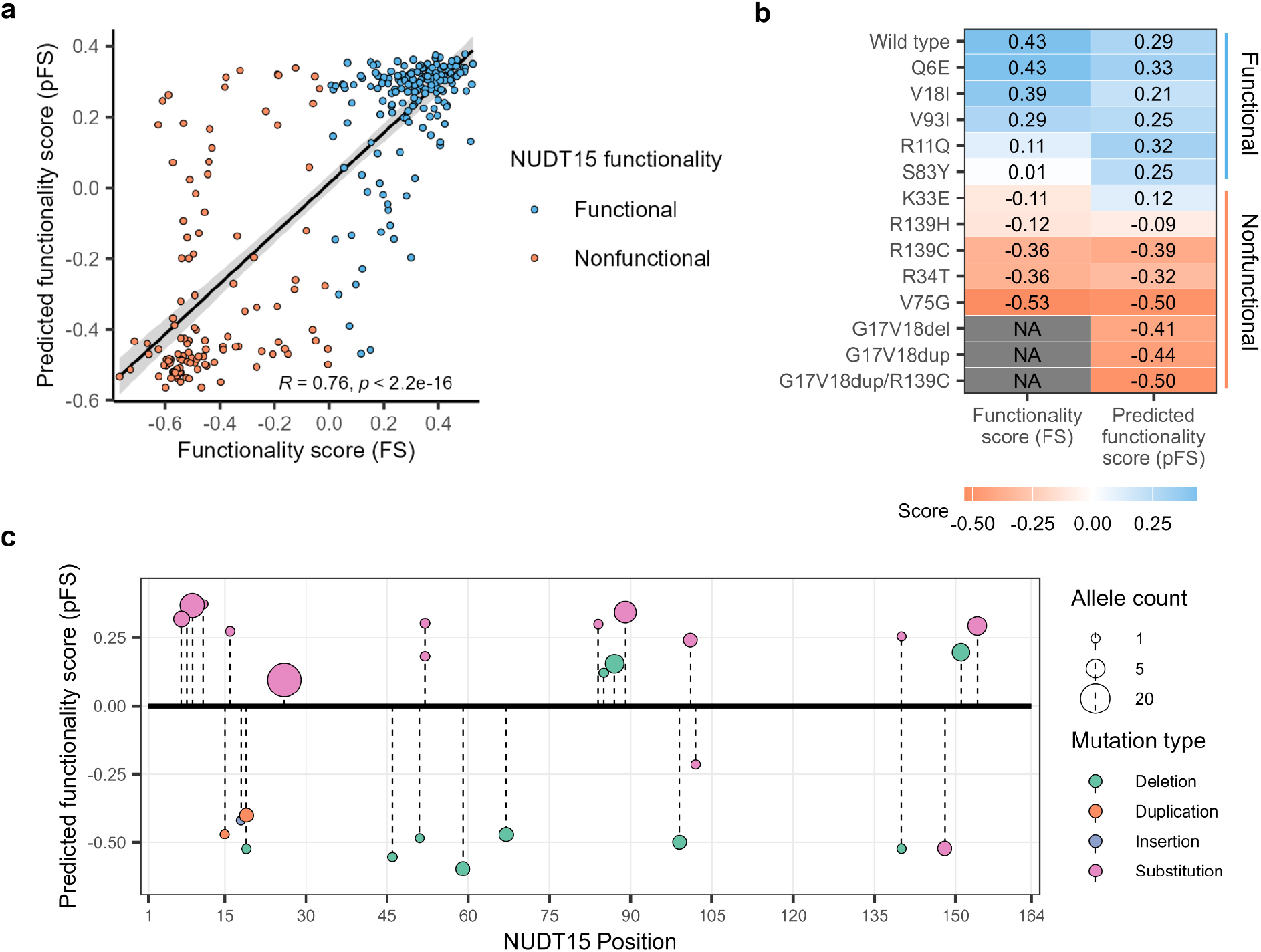
Fine-tuned ESM-2 correlates NUDT15 sequence variation with thiopurine cytotoxicity risk. **a**, pFS values (y-axis) compared to measured FS values (x-axis) for validation sequences are shown for our ESM-2 model fine-tuned on NUDT15 variant data. Negative FS values are in orange and positive FS values are in blue. Line of best fit with 95% confidence interval and Spearman correlation coefficient (R) are also shown. **b**, A table of FS (left) and pFS (right) values for clinically characterized NUDT15 variants is shown, with negative values in orange, positive values in blue, and missing values in grey. NUDT15 variant functionality as determined by thiopurine sensitivity in patients is shown on the right, with the blue bracket denoting benign functional variants and the orange bracket denoting nonfunctional variants that increased thiopurine cytotoxicity. **c**, pFS values (y-axis) for clinically uncharacterized genomAD variants lacking an FS value are shown along the NUDT15 sequence (x-axis). Variants are colored by mutation type and sized by allele count.

We first collected any clinically characterized NUDT15 variants that were benign or associated with thiopurine cytotoxicity^21–23^. This yielded 14 variants, 11 of which had a corresponding FS value from the Suiter *et al*. assay. All variants were scored by our fine-tuned modeling and compared against the FS values and clinical observations (**Fig. 4b**). All benign mutants and cytotoxic missense substitutions were correctly scored by the Suiter *et al*. assay, while our fine-tuned model performed similarly except for one cytotoxic variant (K33E) which was incorrectly scored as benign. Three NUDT15 variants with cytotoxic insertion/deletion mutations were not scored in the Suiter *et al*. assay because only single substitution mutations were tested. In contrast, our fine-tuned model was able to successfully score all three variants as nonfunctional. The Genome Aggregation Database^28^ (genomAD) was searched for additional uncharacterized missense variants of the NUDT15 gene. This returned 29 clinically uncharacterized variants which lacked a corresponding FS value from the Suiter *et al*. assay. These genomAD variants were screened by our fine-tuned model, which scored most in-frame deletion, duplication, and insertion mutants as nonfunctional and most substitution mutants as functional (**Fig 4c**.) Further testing would be necessary to determine if these predictions are accurate for patients possessing such NUDT15 variants.

We demonstrate that fine-tuned PLMs trained on directed evolution data can be used to phenotype previously unseen naturally-evolved genotypic variants. *Pik-1* binding to Avr-PikC appears to be rare in rice, with only two alleles identified recently in wild rice varieties exhibiting strong binding to the ligand^29,30^. The diversity of *Pik-1* HMA domain variants we detected within the 3k RGP dataset supports previous observations that the selective pressure imparted by *M. oryzae* is encouraging diversification of the *Pik-1* HMA domain^31^. The directed evolution methodology we implemented mimics this selective pressure, which ESM-2 can learn from to accurately correlate naturally-occurring sequence variation with changes in ligand binding. Using our fine-tuned models, we identified two *Pik-1* HMA domain variants from the 3k RGP, VK and SHZ-2, which exhibit enhanced binding to Avr-PikC relative to Pikh-1 *in vitro*. Interestingly, the SHZ-2 rice cultivar has been used as a source of blast resistance in current breeding programs without recognition of the potential strength of its Pik-1 allele^32,33^. Whether the improved ligand binding we observed translates into a robust activation of immunity against Avr-PikC *in planta*, and partially contributes to the strong blast resistance of SHZ-2, remains to be tested. Overall, obtaining receptor candidates in this manner has the potential to vastly accelerate the process of testing and developing resilient rice varieties needed by growers around the world.

Transformer-based models have previously shown state-of-the-art performance in correlating genetic variations with phenotypes of interest^34^. Our approach with PLMs further highlights the power of transformers for genotype-to-phenotype analyses. Notably, our modeling accurately predicted enhanced Avr-PikC binding to *Pik-1* variants possessing an insertion in the HMA domain although the training data used for our fine-tuning contained no variants with insertions or deletions. This performance was recapitulated in our modeling of NUDT15 functionality, which accurately predicted the negative impacts of sequence insertions/deletions on thiopurine cytotoxicity risk even after training on a sequence dataset which lacked insertions or deletions. This accuracy indicates that fine-tuned PLMs can be used to effectively gauge the impact of unusual or previously unseen genotypic variations on phenotypes of interest. Ultimately, our approach helped identify immune receptor variants in rice that exhibit rare ligand recognition properties. We also show that the same methodology could be applied toward the prediction of other phenotypes of interest, such as patient drug sensitivity risk. Utilizing genotype-to-phenotype approaches like these will be an increasingly important step towards fully utilizing the wealth of information found in large genomic datasets.

## Acknowledgments

This project was supported by the University of California Davis Flow Cytometry Shared Resource Laboratory with technical assistance from Bridget McLaughlin, Jonathan Van Dyke and Ashley Karajeh.

## Funding

Life Sciences Research Foundation – Simons Foundation (E.Y.R.)

National Institutes of Health MIRA 1R35GM148173 (P.C.R.)

National Institutes of Health grant P30 CA093373 (Flow Cytometry Shared Resource)

National Institutes of Health NCRR C06-RR1208 (Flow Cytometry Shared Resource)

National Institutes of Health S10 OD018223 (Flow Cytometry Shared Resource)

National Institutes of Health S10 RR 026825 (Flow Cytometry Shared Resource)

James B. Pendleton Charitable Trust (Flow Cytometry Shared Resource)

Partially supported by the Joint BioEnergy Institute, U.S. Department of Energy, Office of Science, Biological and Environmental Research Program under Award Number DEAC02-05CH11231 (P.C.R.)

## Supplementary Materials

Methods

Table S1 and S2

