## Supplementary Materials for "Combining Directed Evolution with Machine Learning Enables Accurate Genotype-to-Phenotype Predictions"

**The PDF file includes:**
Methods
Table S1 and S2

**Methods:**

*Directed evolution of the Pik-1 ligand binding domain*

Directed evolution of the Pikh-1 HMA ligand binding domain against Avr-PikC and Avr-PikF was performed as previously described[19]. The 1 µM Avr-PikC and 1 µM Avr-PikF post-sort YSD libraries, as well as the initial starting YSD library, were analyzed with next-generation sequencing to survey the change in receptor populations following FACS selection.

*Scoring Pik-1 variant fitness for Avr-Pik ligand binding*

Next-generation sequencing reads from each YSD library were translated and trimmed to isolate the displayed HMA domains. Variants containing stop codons or insertions/deletions were removed, and the read counts corresponding to each HMA domain variant were quantified. An "enrichment score" was calculated for each variant based on the change in sequence abundance following FACS selection. This score was calculated by taking the log of the ratio of count-normalized post-sort reads to count-normalized starting library reads. After scoring, any sequence variants with fewer than 10 reads in both the starting and post-sort libraries were filtered out of the dataset to ensure variants used for training had high-confidence changes in abundance.

*Modeling Pik-1 variant ligand binding fitness*

Enrichment data for Avr-PikC and Avr-PikF binding was split 0.95/0.05 (training/validation). The HuggingFace Transformers[35] library was used to fine-tune the ESM-2 T6_8M_UR50D base model over 20 epochs to predict pES values based on input training sequences. The fine-tuned model weights which yielded the highest Spearman ranked correlation coefficient R value on validation data predictions were chosen for use in downstream predictions. The embeddings for each sequence in the training/validation sets were extracted from the ESM-2 T6_8M_UR50D base model and averaged across all amino acid positions. These averaged embeddings were used to train a gradient-boosted decision tree in CatBoost[36] (v1.2.7), a cross-validated ElasticNet regression model, and a support vector regression (SVR) model. Cross-validated ElasticNet and SVR models were made with the scikit-learn library (v1.5.2) using default parameters. These ESM-2 sequence embeddings were also clustered using tSNE to manually split the data by latent space into training and validation sets (0.95/0.05). Models were trained on this latent space split data to determine which method was most effective at predicting the binding behavior of out-of-distribution validation sequences. All models were compared by calculating the Spearman R and RMSE of validation data predictions.

*Extracting Pik-1 variants from the 3,000 Rice Genomes Project dataset*

BWA-MEM[37] was used to align short read sequencing data from the 3k RGP against the N22 reference genome (GCA_001952365.3). Samples with full read coverage over the *Pik-1* HMA domain were analyzed with the FreeBayes variant calling pipeline[38].

*G2P analysis of naturally-evolved Pik-1 variant ligand binding*

*Pik-1* HMA domain variants detected in the 3k RGP were input into our Avr-PikC and Avr-PikF fine-tuned ESM-2 models to get pES values for each sequence. The HMA domain variants of interest were cloned into the pCTcon2 YSD plasmid and expressed in EBY100 yeast. Cells were stained for binding against 1 µM Avr-PikA or 1 µM Avr-PikC as previously described[19]. Receptor expression and ligand binding relative to Pikh-1 was captured with fluorescent microscopy on a Leica SP8 confocal microscope. Receptor expression and binding against 1 µM Avr-PikA or 1 µM Avr-PikC of each variant was quantified via FACS on Becton Dickinson FACS Aria II at the UC Davis Flow Cytometry Shared Resource as previously described[19]. Briefly, 1:100 Myc tag rabbit monoclonal antibody (Cell Signaling 2278) and 1:100 anti-rabbit IgG secondary antibody Alexa Fluor 488 conjugate (Thermo Fisher A-11008) were used to detect receptor expression and 1:100 streptavidin PE conjugate (Thermo Fisher S866) was used to detect ligand binding.

*Modeling and phenotyping NUDT15 variant functionality*
NUDT15 variant functionality scores generated by Suiter *et al.*[21] were adjusted and log-normalized to score all functional variants positively and all nonfunctional variants negatively. Clinically characterized variants of NUDT15 were filtered from this dataset for use in downstream phenotyping. All remaining variant data was split 0.90/0.10 (training/validation) for fine-tuning. A fine-tuned ESM-2 model was trained and selected as previously described. Additional NUDT15 missense variants were extracted from genomAD v4.1.0 and filtered for variants without functional scoring by the Suiter *et al.* assay or known clinical significance. This subset of genomAD variants, as well as the clinically characterized NUDT15 variants, were input into our fine-tuned ESM-2 model to get predicted functional scores for each sequence.

*Data visualization*
Figure 1a and 1b were made in BioRender.com and can be found at https://BioRender.com/o43t244. Plots of the distribution of FACS-measured cell fluorescence were made in FlowJo (v10.10.0), where the same receptor expression threshold was applied to all variants and ligand binding of cells exceeding this threshold was plotted as a histogram. All remaining plots were made with R (v4.2.2)[39] in RStudio[40] (v2023.3.1.446) using ggplot2 (v3.4.3) and ggmsa[41] (v1.3.4). Fluorescent microscopy images were cropped and adjusted with the same parameters applied to each image in ImageJ[42] (v2.9.0).

*Data/code availability*
Scripts, source files, and plots generated for this study are available on https://github.com/alexanderjhoward/Pik1-DE-FT.

**Table S1: Directed evolution data is most effectively modeled by fine-tuned ESM-2.** Several models were trained on the same *Pik-1* training/validation data for our Avr-PikC and Avr-PikF directed evolution sorts. Shown are the Spearman R values and RMSE values for the validation sequence predictions made with each modeling approach, with all but the fine-tuned ESM-2 models trained on ESM-2 sequence embeddings. The largest R-value and lowest RMSE value for each validation dataset is shown in bold.

| Model | Validation Data | Spearman R | RMSE |
|---|---|---|---|
| Fine-tuned ESM-2 | Avr-PikC | **0.868** | **0.267** |
| CatBoost | Avr-PikC | 0.810 | 0.314 |
| SVR | Avr-PikC | 0.703 | 0.310 |
| ElasticNet Regression | Avr-PikC | 0.826 | 0.309 |
| Fine-tuned ESM-2 | Avr-PikF | **0.855** | **0.297** |
| CatBoost | Avr-PikF | 0.770 | 0.406 |
| SVR | Avr-PikF | 0.574 | 0.353 |
| ElasticNet Regression | Avr-PikF | 0.785 | 0.382 |

**Table S2: Fine-tuned ESM-2 best predicts the ligand binding of out-of-distribution validation sequences.**

ESM-2 embeddings from Avr-PikC and Avr-PikF directed evolution sorts were collected and clustered using tSNE to manually split the training and validation data by latent space. All models were then trained on this latent space split training/validation data. Shown are the Spearman R values and RMSE values for the validation sequence predictions made with each modeling approach, with all but the fine-tuned ESM-2 models trained on ESM-2 sequence embeddings. The largest R-value and lowest RMSE value for each validation dataset is shown in bold.

| **Model** | **Validation Data** | **Spearman R** | **RMSE** |
|---|---|---|---|
| Fine-tuned ESM-2 | Avr-PikC | **0.686** | **0.350** |
| CatBoost | Avr-PikC | 0.570 | 0.399 |
| SVR | Avr-PikC | 0.362 | 0.447 |
| ElasticNet Regression | Avr-PikC | 0.479 | 0.438 |
| Fine-tuned ESM-2 | Avr-PikF | **0.715** | **0.325** |
| CatBoost | Avr-PikF | 0.517 | 0.462 |
| SVR | Avr-PikF | 0.513 | 0.500 |
| ElasticNet Regression | Avr-PikF | 0.518 | 0.505 |
